# scPNMF: sparse gene encoding of single cells to facilitate gene selection for targeted gene profiling

**DOI:** 10.1101/2021.02.09.430550

**Authors:** Dongyuan Song, Kexin Aileen Li, Zachary Hemminger, Roy Wollman, Jingyi Jessica Li

## Abstract

Single-cell RNA sequencing (scRNA-seq) captures whole transcriptome information of individual cells. While scRNA-seq measures thousands of genes, researchers are often interested in only dozens to hundreds of genes for a closer study. Then a question is how to select those informative genes from scRNA-seq data. Moreover, single-cell targeted gene profiling technologies are gaining popularity for their low costs, high sensitivity, and extra (e.g., spatial) information; however, they typically can only measure up to a few hundred genes. Then another challenging question is how to select genes for targeted gene profiling based on existing scRNA-seq data. Here we develop the single-cell Projective Non-negative Matrix Factorization (scPNMF) method to select informative genes from scRNA-seq data in an unsupervised way. Compared with existing gene selection methods, scPNMF has two advantages. First, its selected informative genes can better distinguish cell types. Second, it enables the alignment of new targeted gene profiling data with reference data in a low-dimensional space to facilitate the prediction of cell types in the new data. Technically, scPNMF modifies the PNMF algorithm for gene selection by changing the initialization and adding a basis selection step, which selects informative bases to distinguish cell types. We demonstrate that scPNMF outperforms the state-of-the-art gene selection methods on diverse scRNA-seq datasets. Moreover, we show that scPNMF can guide the design of targeted gene profiling experiments and cell-type annotation on targeted gene profiling data.

## 1 Introduction

The recent development of single-cell RNA sequencing (scRNA-seq) technologies provides unprecedented opportunities to decipher transcriptome heterogeneity among individual cells [1–3]. A typical scRNA-seq dataset contains thousands to tens of thousands of genes; however, a subset of genes, which we call *informative genes*, are usually sufficient for representing the underlying biological variations of cells in the dataset for two reasons. First, variations of many genes are not related to the biological variations of interest. For instance, fluctuations in the expression levels of housekeeping genes are irrelevant to cell types [4, 5]. Second, many genes have strongly correlated expression levels, suggesting that one gene may represent a group of genes without much loss of information [6]. Therefore, for scRNA-seq data analysis, informative gene selection has three advantages: (1) enhancing biological signals by removing unwanted technical variations, (2) improving the interpretability of analysis results by focusing on informative genes, and (3) reducing the number of genes to save computational resources.

Besides scRNA-seq data analysis, informative gene selection is also crucial for designing single-cell targeted gene profiling experiments, which we define to include all technologies that measure only a specific sets of genes’ expression levels in individual cells. Unlike scRNA-seq, targeted gene profiling requires a limited number (often no more than hundreds) of genes to be specified before sequencing. Examples of targeted gene profiling include spatial technologies (e.g., smFISH [7] and MERFISH [8]) and non-spatial technologies (e.g., BART-Seq [9], HyPR-seq [10] and 10x-Genomics Targeted Gene Expression). Compared with scRNA-seq, targeted gene profiling technologies have advantages such as capturing spatial information (by smFISH and MERFISH), having a lower cost per cell (by BART-Seq), and exhibiting a higher sensitivity for detecting lowly expressed genes (by HyPR-seq). However, it remains an open and challenging question to optimize the gene selection for targeted gene profiling under a gene number limitation.

Given the importance of informative gene selection, researchers have developed many gene selection methods for scRNA-seq data. Most existing methods select genes based on the relationship between per-gene expression means and per-gene expression variances (with the mean and variance of each gene calculated across cells). Popular example methods include variance stabilization transformation (vst) [11] and mean-variance plot (mvp) in the R package Seurat [12], as well as modelGeneVar in the R package scran [13]. These methods select highly variable genes that have large expression variances in relation to their expression means. Other methods use various metrics of gene importance instead of the per-gene expression variance. For example, M3Drop selects the genes that have zero expression levels in many cells [14]; GiniClust selects the genes with large Gini indices of expression levels [15]; SCMarker selects the genes that have expression levels bi/multi-modally distributed and are co-expressed or mutually-exclusively expressed with some other genes [16]. A common limitation of these existing methods is that they are all designed to select a relatively large number of genes; thus, their performance in selecting a small number of genes remains unclear. For instance, in Seurat, the default gene number is 2000; SCMarker selects 700-900 genes in its exemplar applications [16]. All these gene numbers are much greater than 200, the maximum gene number allowed by multiple targeted gene profiling technologies. Therefore, existing gene selection methods may not be suitable for selecting genes for targeted gene profiling. Another drawback of these methods is that their selected genes lack functional interpretability; that is, their selected genes are not categorized as functional gene groups.

In addition to these gene selection methods, linear dimensionality reduction methods, such as principal component analysis (PCA) and non-negative matrix factorization (NMF), can also used for gene selection. Specifically, genes can be selected based on their contributions to the projected low dimensions found by PCA or NMF [17–19]. Although many variants of PCA and NMF algorithms have been developed for scRNA-seq data analysis, they are not designed for gene selection [20–26].

Here we propose an unsupervised method scPNMF to simultaneously select informative genes and project scRNA-seq data onto an interpretable low-dimensional space. Leveraging the Projective Non-negative Matrix Factorization (PNMF) algorithm [27], scPNMF combines the advantages of PCA and NMF by outputting a non-negative sparse weight matrix that can project cells in a high-dimensional scRNA-seq dataset onto a low-dimensional space. Unlike the weight matrix (a.k.a., loading matrix) found by PCA, the non-negative sparse weight matrix output by scPNMF correspond to bases that each correspond to a group of co-expressed genes. Compared with the original PNMF, a unique feature of scPNMF is basis selection: scPNMF uses correlation screening and multimodality testing to remove the bases that cannot reveal potential cell clusters in the input scRNA-seq dataset. There are two functionalities of scPNMF: (1) given a pre-specified gene number and a scRNA-seq dataset, scPNMF selects informative genes based on its weight matrix; (2) given a targeted gene profiling dataset containing the informative genes, scPNMF projects this dataset onto the same low-dimensional space of a reference scRNA-seq dataset containing cell type labels, thus enabling cell type annotation on the targeted gene profiling dataset. Comprehensive benchmark shows that scPNMF outperforms existing gene selection methods in two aspects. First, the informative genes selected by scPNMF lead to the most accurate cell clustering. Second, the informative genes and weight matrix of scPNMF lead to the best cell type prediction accuracy for targeted gene profiling data. Therefore, scPNMF is a powerful gene selection method that can guide the experimental design and data analysis of single-cell targeted gene profiling.

## 2 Methods

The core of scPNMF is to learn a low-dimensional embedding of cells so that the bases of the low-dimensional space correspond to sparse and mutually exclusive gene groups, and that genes in each group are co-expressed and thus functionally related. Fig.1 illustrates the workflow of scPNMF. The input of scPNMF is a log-transformed gene-by-cell count matrix measured by scRNA-seq. There are two main steps in scPNMF: (I) it learns a low-dimensional sparse weight matrix by PNMF; (II) it selects bases in the weight matrix based on functional annotations (optional), correlation screening, and multimodality testing to remove uninformative bases that cannot distinguish cell types. The output of scPNMF includes (1) the selected weight matrix, a sparse and mutually exclusive encoding of genes as new, low dimensions, and (2) the score matrix containing embeddings of input cells in the low dimensions. The selected weight matrix has two main applications: extracting informative gene for downstream analyses, such as cell clustering and new marker gene identification, and projecting new targeted gene profiling data for data integration and cell type annotation.

**Figure 1:**
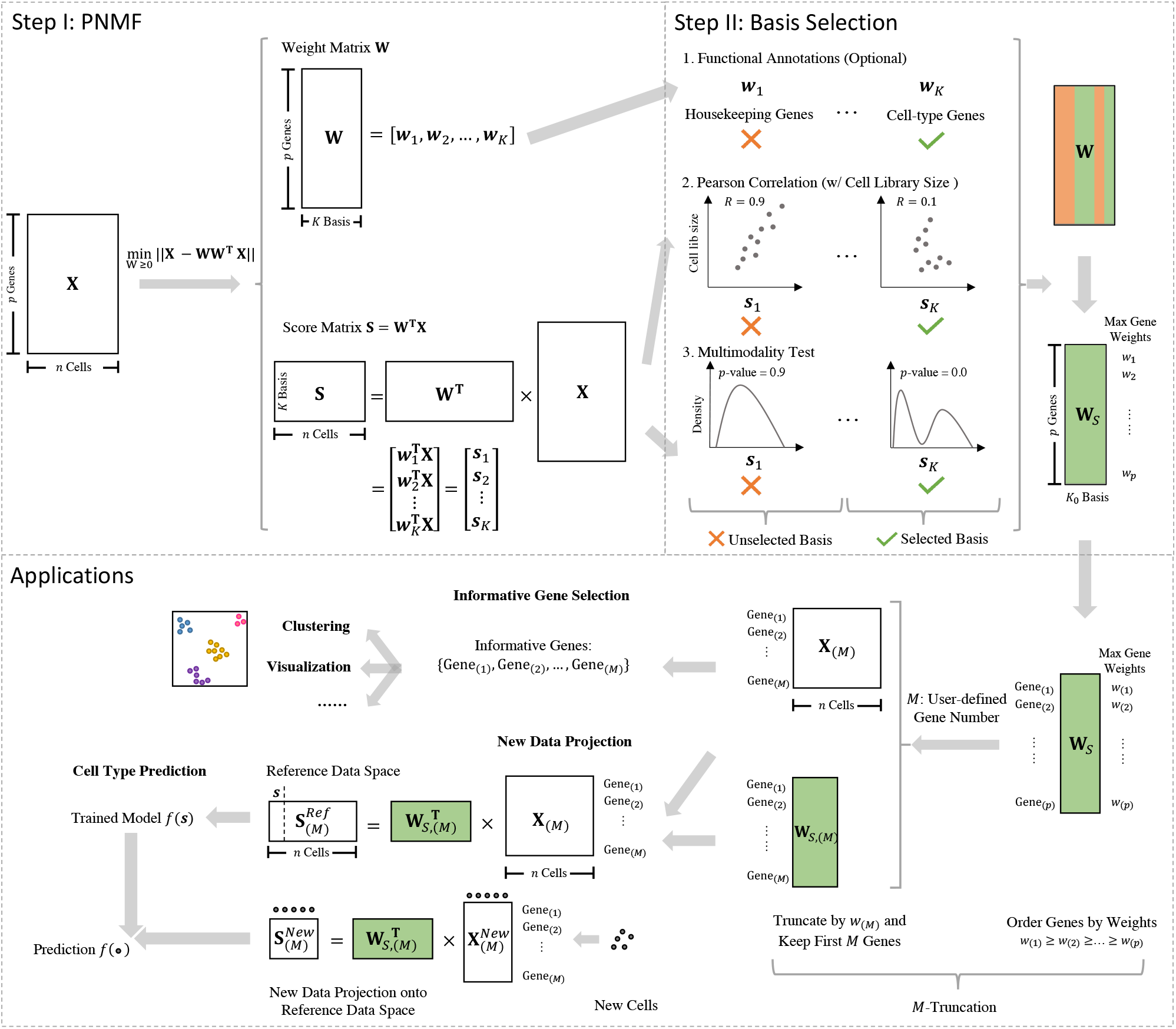
An overview of scPNMF. Taking a log-transformed gene-by-cell count matrix as the input, scPNMF first learns a low-dimensional sparse weight matrix **W** and a low-dimensional cell embedding matrix **S**. Second, it remove the bases irrelevant to cell type variations by examining bases’ functional annotations (optional), Pearson correlations with cell library sizes, and multimodality. Given a user-defined gene number *M*, scPNMF performs *M*-truncation to facilitate two main applications: (1) selecting the desired number of informative genes; (2) projecting new targeted gene profiling data onto the low-dimensional space defined by reference scRNA-seq data. The details are in the “Methods” section.

### 2.1 scPNMF step I: PNMF

In this section, we review the PNMF algorithm [27, 28] as the foundation of scPNMF. We first compare the formulation of PNMF with that of principal component analysis (PCA) and non-negative matrix factorization (NMF), and we show that PNMF has the advantages of both PCA and NMF so that it can be a useful tool for scRNA-seq data analysis. Next, we introduce our PNMF implementation.

Given a log-transformed count matrix 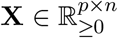, whose *p* rows correspond to genes and whose *n* columns represent cells, and a positive integer *K ≤ p*, PNMF aims to find a *K*-dimensional space, whose dimensions correspond to non-negative, sparse and mutually exclusive linear combinations of the *p* genes, so that projecting the *n* cells onto the *K*-dimensional space does not cause much information loss (i.e., projecting the *K*-dimensional embeddings of the *n* cells back to the original *p*-dimensional space can largely restore the original *n* cells). PNMF tackles this task by solving the optimization problem:

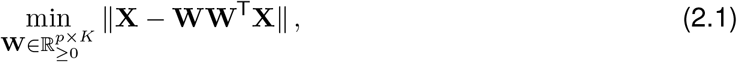

where *‖ · ‖* denotes the Frobenius matrix norm. The solution **W** is referred to as a *weight matrix*. Each column of **W** is a *basis*, whose *p* entries are the weights of the *p* genes. PNMF requires all weights to be non-negative, leading to a sparse **W** with most weights as zeros.

PCA is similar to PNMF but does not require all weights to be non-negative. We can write the optimization problem of PCA as

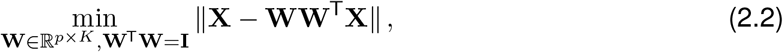

whose solution **W** is also a weight matrix but not sparse, and **W** is often referred to as the loading matrix.

A common property of PNMF and PCA is that the transpose of their weight matrix, **W**^T^ *∈* ℝ^*K×p*^, can be used to project a new cell with *p* gene measurements, **x** *∈* ℝ^*p*^, onto the *K*-dimensional space as **W**^T^**x**.

In contrast to PMNF and PCA, NMF finds two non-negative matrices **W** and **H** so that their product approximates the original matrix **X**. NMF solves the optimization problem:

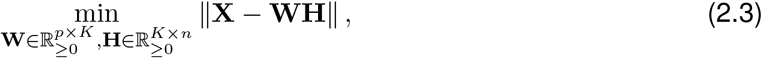

whose solution **W** still has *K* columns representing bases, and **H** has *n* columns as *K*-dimensional embeddings of the *n* cells. Due to the non-negative constraint on **W** and **H, W** is a sparse matrix [29]. However, the transpose **W**^T^ cannot be used as a projection matrix from the original *p*-dimensional space to a *K*-dimensional space. The reason is that, if **W**^T^ is a projection matrix, then by the definition of **H** we have **W**^T^**X** = **H**, which would converts the objective function (2.3) of NMF to the objective function (2.1) of PNMF. In other words, PNMF is a constrained version of NMF by requiring **W**^T^ to be a projection matrix. Hence, PNMF inherits the property of NMF by having non-negative, sparse bases that are mostly mutually exclusive (i.e., different bases correspond to different gene groups). Moreover, based on the similarities of the objective functions of PNMF (2.1) and PCA (2.2), we can see that PNMF also resembles PCA by finding a weight matrix whose transpose can serve as a projection matrix and whose bases are largely orthogonal to each other. Table 1 summarizes the properties of PNMF, PCA, and NMF.

**Table 1:** Comparison of the properties of PNMF, PCA and NMF

|  | Optimization Problem | Non-negativity | Sparsity | Mutually Exclusiveness | New Data Projection |
| --- | --- | --- | --- | --- | --- |
| PNMF | $\min_{\mathbf{W}} \ \mathbf{X} - \mathbf{WW}^T \mathbf{X}\ \text{ s.t. } \mathbf{W} \geq 0$ | Yes | Very high | Very high | Yes |
| PCA | $\min_{\mathbf{W}} \ \mathbf{X} - \mathbf{WW}^T \mathbf{X}\ \text{ s.t. } \mathbf{W}^T \mathbf{W} = \mathbf{I}$ | No | Low | Low | Yes |
| NMF | $\min_{\mathbf{W}, \mathbf{H}} \ \mathbf{X} - \mathbf{WH}\ \text{ s.t. } \mathbf{W}, \mathbf{H} \geq 0$ | Yes | High | High | No |

In the context of scRNA-seq data analysis, the above advantages of PNMF lead to an interpretable and useful weight matrix **W**. First, the high sparsity of **W** makes each basis (column) depend on only a small set of genes, which has been defined as a *meta-gene* for NMF [30]. Second, the mutual exclusiveness of **W** makes different bases correspond to different gene sets, easing the interpretation of bases as meta-genes or functional units. Third, the projection matrix **W**^T^ allows the alignment of new data to reference data, thus facilitating cell type annotation on the new data.

Algorithm 1 summarizes the key steps of PNMF implementation in scPNMF. Our implementation mainly follows the two papers that proposed the PNMF algorithm [27, 28], and we change the initialization of **W** to the weight matrix found by PCA, **W**_PCA_, with the absolute value taken on every entry. Our initialization is motivated by the desired orthogonality of bases (i.e., columns of **W**).

With the weight matrix 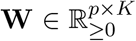 learned by PNMF, we obtain the *score matrix* 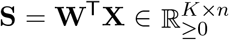, whose *K* rows correspond to the bases and whose *n* columns represent the cells. Specifically, the *j*-th column of **S** is the *K*-dimensional embedding of the *j*-th cell; the *k*-th row of **S**,

#### Algorithm 1 seudocode of PNMF implementation in scPNMF

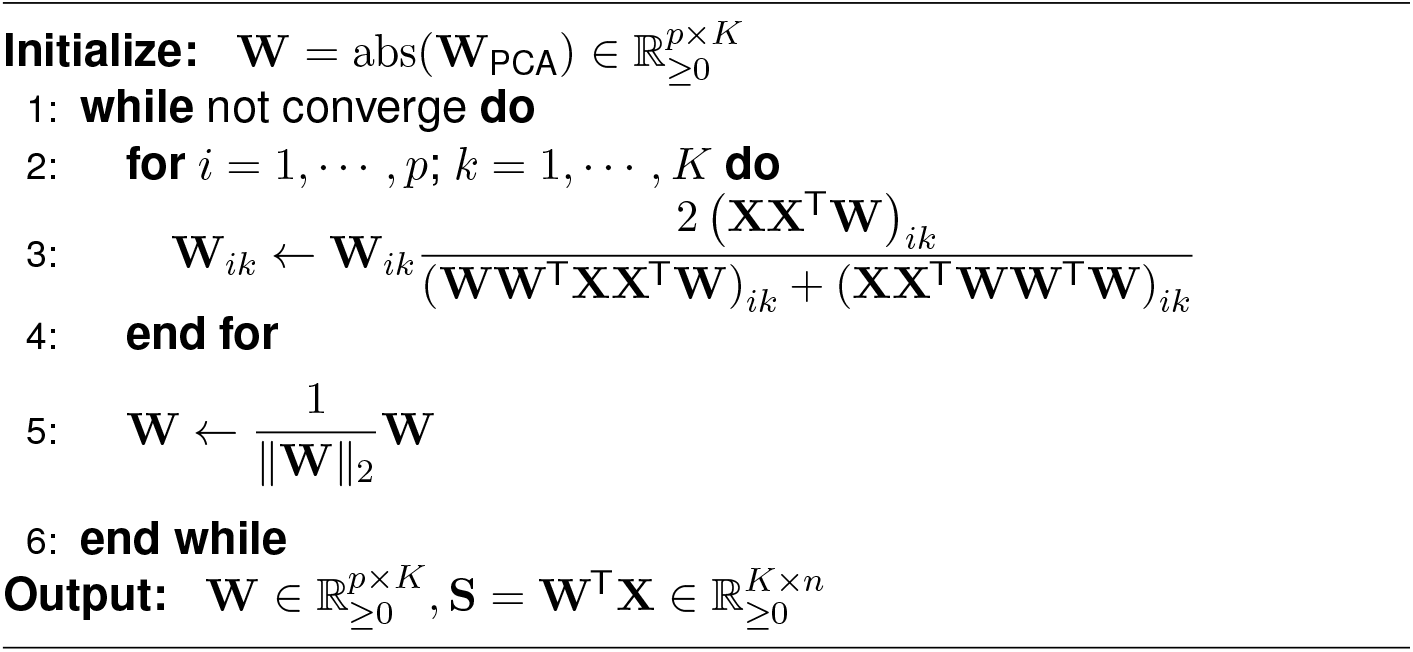

denoted by 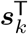, contains the *scores* (i.e., coordinates) of all *n* cells in the *k*-th basis:

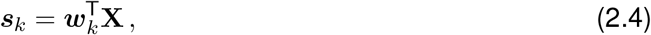

where ***w***_*k*_ is the *k*-th column of **W**, *k* = 1, …, *K*.

The low rank *K* needs to be pre-specified in PNMF, same as in PCA and NMF, A larger *K* preserves more information in **X** but also removes less noise (technical variation of cells that is not of biological interest), impedes the interpretation of **W** (more bases are more difficult to interpret), and increases the computational burden. To choose *K* in a data-driven way, we propose an orthogonality measure, which shows that *K* = 20 is a reasonable choice for multiple scRNA-seq datasets (Section S1.1).

### 2.2 scPNMF step II: basis selection

The second key step of scPNMF is to select informative bases among the *K* bases found by PNMF (i.e., columns of **W** and rows of **S**) to remove unwanted variations of cells (e.g., variations irrelevant to cell types). The columns of **W** enjoy high sparsity and mutual exclusiveness; that is, each column contains positive weights corresponding to a unique small set of genes, so it is expected to reflect a certain biological function. However, some biological functions may not be relevant to the cell heterogeneity of interest, e.g., cell type composition. Motivated by this, we propose three strategies for selecting informative bases (columns of **W** and rows of **S**): functional annotations (optional), correlations with cell library sizes, and tests of multimodality.

#### 2.2.1 Strategy 1: examine bases by functional annotations (optional)

The first, optional strategy is to annotate the biological function(s) of each basis in the weight matrix. For example, scPNMF may apply gene ontology (GO) analysis to the top 10% genes with the highest weights in each basis (column of **W**) and record the enriched GO terms as the basis’ functional annotation. Then, users with prior knowledge can interpret the functional annotation on each basis and decide whether or not to remove the basis. For example, if the goal is to delineate cell types in scRNA-seq data, a basis corresponding to cell-cycle genes should be removed because they would obscure the distinction of cell types.

However, it is worth noting that filtering bases by biological annotations is optional in scPNMF. Conservative users can keep all *K* bases output by PNMF and directly use data-driven basis selection (Section 2.2.2). For our results in this paper, scPNMF removes the bases corresponding to well-known housekeeping genes (Section S2).

#### 2.2.2 Data-driven strategies

##### 2.2.2.1 Strategy 2: examine bases by correlations with cell library sizes

Note that the input of scPNMF is a log-transformed unnormalized count matrix for users’ convenience. Hence, scPNMF does not adjust for cell library sizes in the computation of **W** and **S** in step I. Given that the variance of cell library sizes contributes to unwanted variations of cells [11], it is necessary to remove the bases whose corresponding rows in **S** are strongly correlated with cell library sizes.

We use the total log-transformed counts to approximate the library size of each cell, and we calculate the Pearson correlation between each ***s***_*k*_ and the library sizes of *n* cells. The strategy is to retain the bases whose Pearson correlations are under a pre-defined threshold, which we set to 0.7 based on empirical observations (Section S1.2).

##### 2.2.2.2 Strategy 3: examine bases by multimodality tests

Another data-driven strategy is to retain the bases whose corresponding scores are multi-modally distributed. If a basis’ score vector (row in **S**) contains *n* scores with a multimodality pattern, then it is likely to distinguish cell types and should be retained. To implement this strategy, we use the ACR test [31] to check the multimodality of each basis’ score vector. The null hypothesis is that the score vector contains *n* scores sampled from a unimodal distribution, and the alternative hypothesis is that the distribution has more than one mode. After performing multiple multimodality tests, one per basis, we use the Benjamini-Hochberg procedure to set a p-value threshold by controlling the false discovery rate under 1%. The bases whose p-values are under this threshold will be retained.

In summary, scPNMF step II allows users to use strategy 1 to filter out uninformative bases based on functional annotations if available; then it implements data-driven strategies 2 and 3 to further remove bases that have strong correlations with cell library sizes and exhibit unimodality patterns. The retained bases will have their corresponding columns in **W** selected and stacked into the *selected weight matrix* 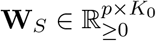, where *K*_0_ is the number of selected bases.

### 2.3 Applications of scPNMF output: informative gene selection and new data projection

The selected weight matrix **W**_*S*_ output by scPNMF has two main applications: selection of a desired number of informative genes and projection of new targeted gene profiling data onto the low-dimensional space defined by **W**_*S*_. Given a gene number *M* (e.g., 200), scPNMF uses *M* - truncation, a step to select *M* rows in **W**_*S*_, resulting in *M* informative genes and a *truncated, selected weight matrix* 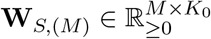 for new data projection.

#### 2.3.1 M-truncation and informative gene selection

We denote the desired number of informative genes by *M ∈* N, with *M ≤* # of non-zero rows in **W**_*S*_. *M*-truncation has three steps.

1. For each gene *i*, calculate its largest weight *w*_*i*_ across bases in **W**_*S*_:

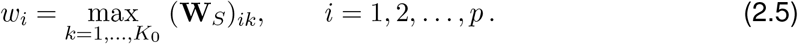
2. Order genes by their maximum weights *w*_(1)_ *≥ w*_(2)_ *≥ · · · ≥ w*_(*p*)_ and set the truncation threshold as *w*_(*M*)_. Identify the first *M* genes as *informative genes*.
3. Construct the truncated, selected weight matrix **W**_*S*,(*M*)_:
  1. Truncate the selected weight matrix **W**_*S*_ by setting all (**W**_*S*_)_*ik*_ *< w*_(*M*)_ to be 0;
  2. Keep the *M* rows with non-zero entries; stack them by row into **W**_*S*,(*M*)_ based on the order of the informative genes.

In short, scPNMF selects informative genes based on their maximum weights in the selected bases. The rationale is that a gene’s maximum weight reflects the gene’s contribution to the establishment of the *K*_0_-dimensional space, which preserves the *n* cells’ biological variations of interest. Hence, genes with larger maximum weights are more informative in the sense of encoding cells’ biological variations. An important application of informative gene selection is to guide the design of targeted gene profiling experiments.

#### 2.3.2 New data projection

Given the selected *M* informative genes, once new cells are measured by targeted gene profiling on these genes, **W**_*S*,(*M*)_ can be used to project the new cells onto the *K*_0_-dimensional space where the cells in the input scRNA-seq data are embedded in. If the input data has cell type annotations, we refer to the input data as *reference data*, then we can predict the new cells’ types from the types of the cells in the reference data. In detail, new data projection has the following steps:

1. Apply scPNMF with *M*-truncation to input, reference data 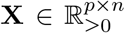with *n* cells to obtain the truncated, selected weight matrix **W**_*S*,(*M*)_. Construct 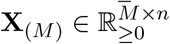 as a submatrix of **X**, with rows corresponding to the rows of **W**_*S*,(*M*)_, i.e., the *M* informative genes. Hence, the *K*_0_-dimensional embeddings of the *n* cells in the reference data are the columns of

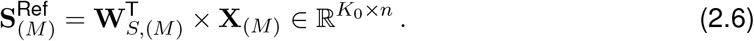
2. Denote the targeted gene profiling data of *n*^*′*^ new cells with *M* informative genes measured by 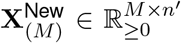. Note that 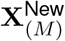 contains log-transformed counts and has rows (genes) corresponding to the rows of **X**_(*M*)_. Project the *n*^*′*^ cells to the *K*_0_-dimensional space by

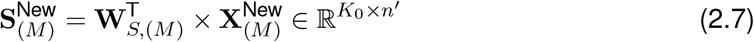
3. (Optional) Normalize 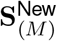 and 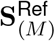 to remove batch effects, if existent, by using a single-cell integration method such as Harmony [32].

Now the *n* reference cells and the *n*^*′*^ new cells are in the same *K*_0_-dimensional space with biological variations preserved. Then a classifier can be trained on the *n* reference cells’ types and 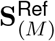 for cell type prediction, and it can be used to predict the *n*^*′*^ cells’ types from 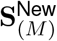

## 3 Results

### 3.1 scPNMF outputs a sparse and functionally interpretable representation of scRNA-seq data

We first demonstrate that scPNMF step I, PNMF, outputs a sparse and functionally interpretable gene encoding of cells. We use the FregGold dataset [33], which consists of three cell types (three human lung adenocarcinoma cell lines), and set the basis number *K* = 5 for demonstration purpose. Both PCA and PNMF learn a weight matrix that can project the original scRNA-seq data onto a 5-dimensional space. Unlike the weight matrix of PCA that has no zero entries, the weight matrix of PNMF is non-negative, highly sparse, containing 42.6% of entries as zeros, and has bases that are largely mutually exclusive (i.e., non-zero entries in different columns correspond to different rows/genes) (Fig. 2a). GO enrichment analysis shows that high weight genes in each PNMF basis are enriched with conceptually-similar GO terms, and high weight genes in different PNMF bases are enriched with conceptually-different GO terms (Fig. 2b). This result indicates that PNMF bases correspond to gene groups with distinct functions. On the contrary, the PCA bases do not have good functional interpretations: the high weight genes in each PCA basis are not enriched with conceptually-similar GO terms, and different PCA bases share many high weight genes (Fig. S3).

**Figure 2:**
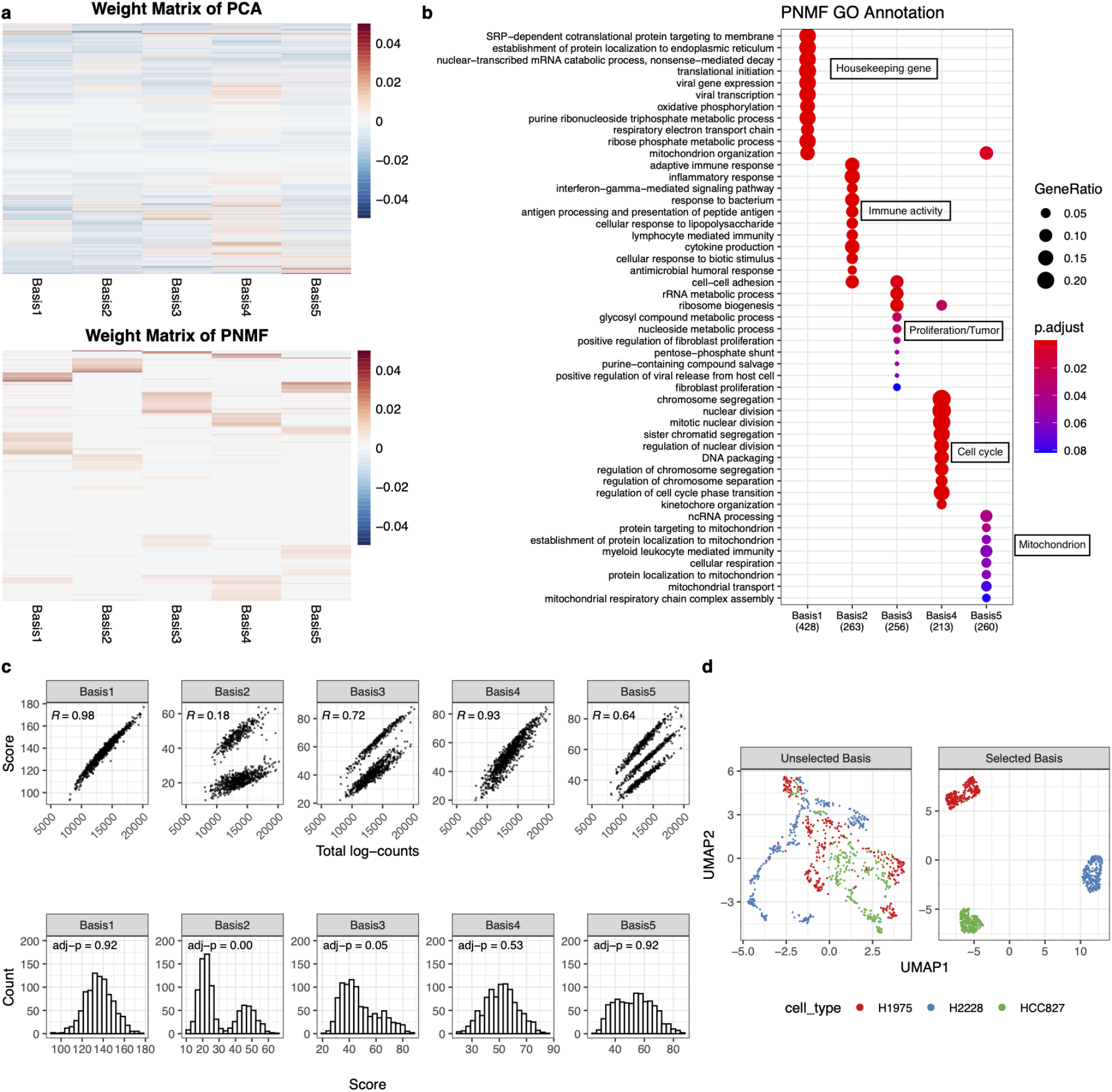
Illustration of the sparse and interpretable projection found by scPNMF. We use the FregGold dataset as an example. (a) Comparison of the weight matrices of PCA and PNMF. Heatmaps visualize the learned weight matrices of PCA (top) and PNMF (bottom), where rows are genes and columns are bases. Red represents positive weights while blue represents negative weights. The rows are ordered by gene-wise hierarchical clustering. Compared to PCA, the weight matrix of PNMF is strictly non-negative, much more sparse and mutually exclusive between bases. (b) GO analysis result of each basis in the weight matrix of PNMF. Texts in black boxes summarize the functions of genes in each basis. The enriched GO terms are almost mutually exclusive, implying that each basis represents a unique gene functional cluster. (c) Statistical tests on each basis in the score matrix of PNMF. Top row: scatter plots of scores and total log-counts (cell library sizes). Each dot represents a cell. Cell scores in bases 1 and 4 are highly correlated with cell library sizes. Bottom row: histograms of cell scores in each basis. Scores in bases 2 and 3 show strong multimodality patterns (adjusted p-value *≤*0.05). (d) UMAP visualizations of cells based on high weight genes in the unselected bases 1 and 4 and those in the selected bases 2, 3, and 5. Genes in the unselected bases completely fail to distinguish the three cell types, while genes in the selected bases lead to a clear separation of the three cell types.

To further analyze the PNMF bases, we list the top 10 high weight genes in each basis (Table S1), from which we identify many well-known genes with important functions. For instance, basis 1 contains classic housekeeping genes, such as *GAPDH* [34] *and ribosomal protein genes (RPS-*) [35]; basis 3 contains well-known tumor-related genes, including *EGFR* [36] *and CDK4* [37]. *In particular, the cells of the HCC827 cell line (one of the three cell types) have overall high scores in basis 3 (Fig. S4), a reasonable result because the HCC827 cell line contains an EGFR* activating mutation [38]. In summary, scPNMF step I outputs bases representing sparse and functionally interpretable gene sets.

### 3.2 Basis selection is an essential step in scPNMF

Here we explain why basis selection is an essential step in scPNMF. In the last section, we show that each PNMF basis of the FregGold dataset approximately represents one functional gene group. It is well known that housekeeping genes (basis 1) and cell-cycle genes (basis 4) are usually irrelevant to cell type distinctions. However, such biological knowledge is not always available or certain. Therefore, scPNMF mainly relies on the two data-driven strategies: correlations with cell library sizes and multimodality tests (Section 2.2.2) for selecting informative bases.

Fig. 2c visualizes the two strategies: cell scores in bases 1 and 4 are highly correlated with cell library sizes (Pearson correlations *>* 0.9); cell scores in bases 2 and 3 show strong evidence as multi-modally distributed (adjusted p-value *<* 0.05). Hence, strategy 1 will not retain bases 1 and 4, and strategy 2 will not retain bases 1, 4, and 5; together, bases 1 and 4 will be removed, and bases 2, 3, and 5 will be selected. To verify the effectiveness of basis selection, we use UMAP to visualize cells based on the top 50 high weight genes in the unselected bases 1 and 4 vs. those in the selected bases 2, 3, and 5 (Fig. 2d). We observe that the top genes in the unselected bases completely fail to separate the three cell types, while the top genes in the selected bases perfectly distinguish the three cell types. This result strongly supports that basis selection is a necessary step of scPNMF.

### 3.3 scPNMF outperforms state-of-the-art gene-selection methods on diverse scRNA-seq datasets

In this section, we demonstrate scPNMF’s capacity for informative gene selection. We comprehensively benchmark scPNMF against 11 other single cell informative selection methods (Table S2) on seven scRNA-seq datasets (Table S3) using three clustering methods (Louvain clustering, K-means clustering, and hierarchical clustering). For fair benchmarking, the seven scRNA-seq datasets cover both unique molecule identifier (UMI) and non-UMI protocols and include various biological samples. Using the adjusted Rank index (ARI) as the metric of clustering accuracy, we calculate the ARI values of the three clustering methods on each dataset using 100 informative genes selected by each gene selection method, as 100 genes are commonly used in targeted gene profiling.

Fig. 3a shows that scPNMF has overall the highest ARI values across datasets and clustering methods. In particular, scPNMF has the highest average ARI value with each clustering method (Louvain: 0.83; K-means: 0.74; hierarchical clustering: 0.69) and the highest overall average ARI (0.75) across datasets and clustering methods. Note that the mean of the overall average ARI values of all methods except scPNMF is only 0.66.

**Figure 3:**
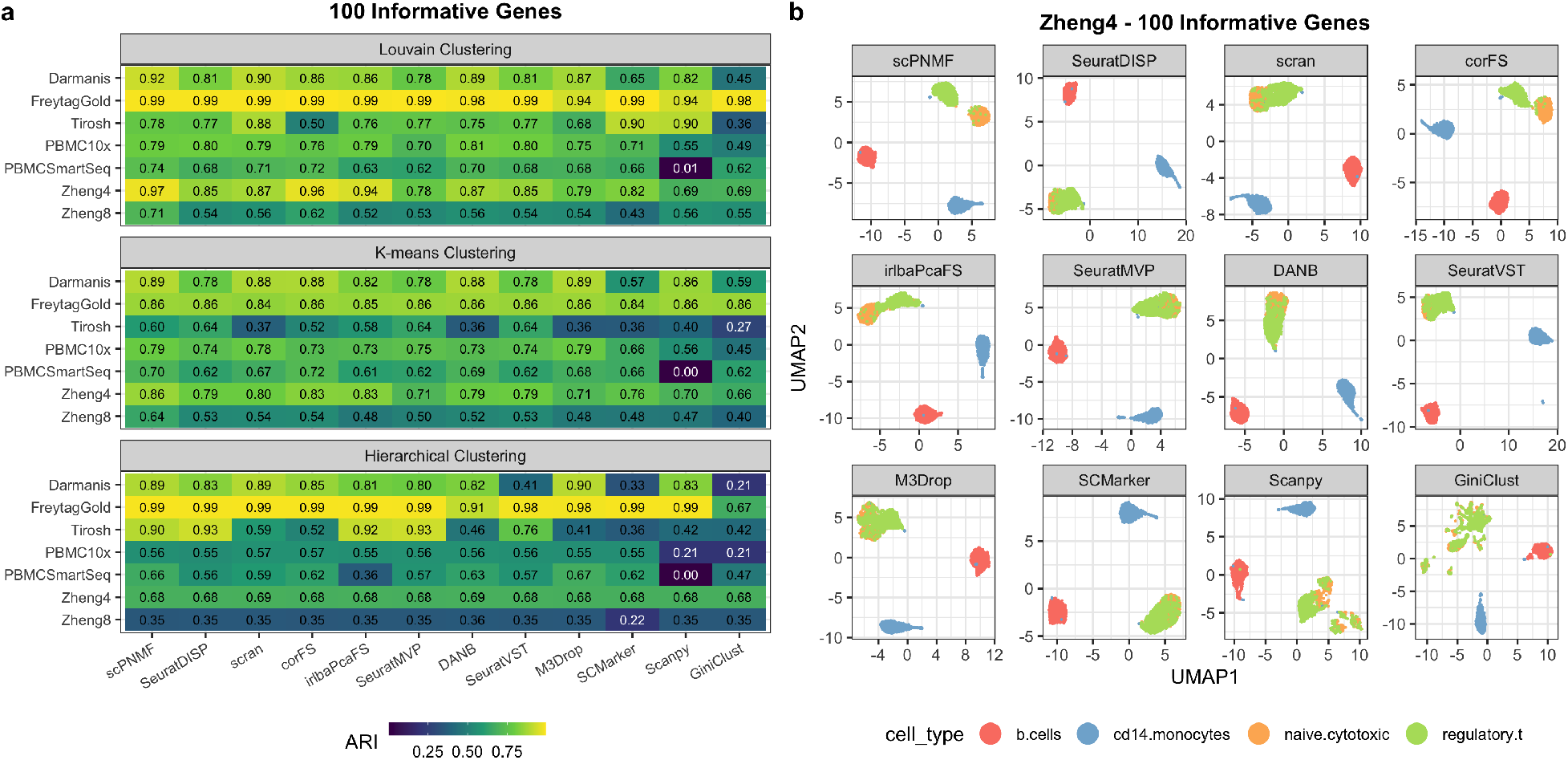
Benchmarking scPNMF against 11 informative gene selection methods on seven scRNA-seq datasets. (a) Clustering accuracies (ARI values) of three clustering methods based on the informative genes selected. Gene selection methods are ordered from left to right by their average ARI across the three clustering methods and the seven datasets. (b) UMAP visualization of cells in the Zheng4 dataset based on 100 informative genes selected by each method. Genes selected by scPNMF lead to a clear separation between naive cytotoxic T cells and regulatory T cells, while the genes selected by others methods do not.

We further show the UMAP visualization of cells in the Zheng4 dataset based on the informative genes selected by each of the 12 gene selection methods (Fig. 3b). Only scPNMF leads to a clear separation of naive cytotoxic T cells and regulatory T cells, while the informative genes selected by other methods except corFS and irlbaPcaFS cannot distinguish the two cell types at all.

We also compare the 12 methods under a varying number of informative genes: 20, 50, 200, and 500, the commonly used gene numbers in targeted gene profiling. We observe that the overall average ARI values of scPNMF are consistently higher than those of other methods, across all informative gene numbers (Fig. S6). Moreover, compared with other methods, scPNMF leads to more stable overall average ARI values under varying numbers of informative genes, indicating its stronger robustness to the gene number constraint of targeted gene profiling. These results strongly support the superior performance of scPNMF as an informative gene selection method.

### 3.4 scPNMF guides targeted gene profiling experimental design and cell-type prediction

In this section, we demonstrate how scPNMF can guide the selection of genes to be measured in a targeted gene profiling experiment, and how scPNMF enables subsequent cell type annotation on the targeted gene profiling data. We design two case studies with paired scRNA-seq reference data and “pseudo” targeted gene profiling data, whose per-cell sequencing depth is higher than that of the corresponding scRNA-seq data.

In the first case study, we use the Zheng8 dataset (measured by the 10x protocol) as the reference dataset. To generate the pseudo targeted gene profiling data, we use a new single-cell gene expression simulator that captures gene correlations, scDesign2 [39], to generate data with a 100-time higher per-cell sequencing depth. In the second case study, we use the PBMC10x dataset (measured by 10x protocol) as the reference dataset, and we use PBMCSmartseq (measured by Smart-Seq2) as the pseudo targeted gene profiling data because Smart-Seq2 has a higher pergene sequencing depth than 10x does. In both case studies, for each gene selection method, the corresponding pseudo targeted gene profiling datasets only contain the *M* informative genes selected by the method.

We benchmark scPNMF against the 11 gene selection methods in terms of cell type prediction on the pseudo targeted gene profiling data. To avoid the bias for a specific classification algorithm, we apply three popular algorithms for cell type prediction: random forest (RF) [40], k-nearest neighbors (KNN) [41], and support vector machine (SVM) [41]. In each case study, we first train each classification algorithm on the low-dimensional embeddings of the reference cells 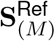 given the *M* = 100 informative genes selected by each gene selection method. Then we apply the trained classifier to the low-dimensional embeddings of the cells in the pseudo targeted gene profiling data 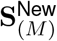. Table 2 shows that scPNMF leads to the highest average prediction accuracy (0.81) across six combinations (two case studies *×* three classification algorithms). Moreover, scPNMF achieves the highest accuracy in each combination except Zheng8 + random forest where it is the second best. These results confirm that scPNMF effectively guides the selection of genes to measure in targeted gene profiling experiments, and it enables accurate cell type annotation on newly generated targeted gene profiling datasets.

**Table 2:**
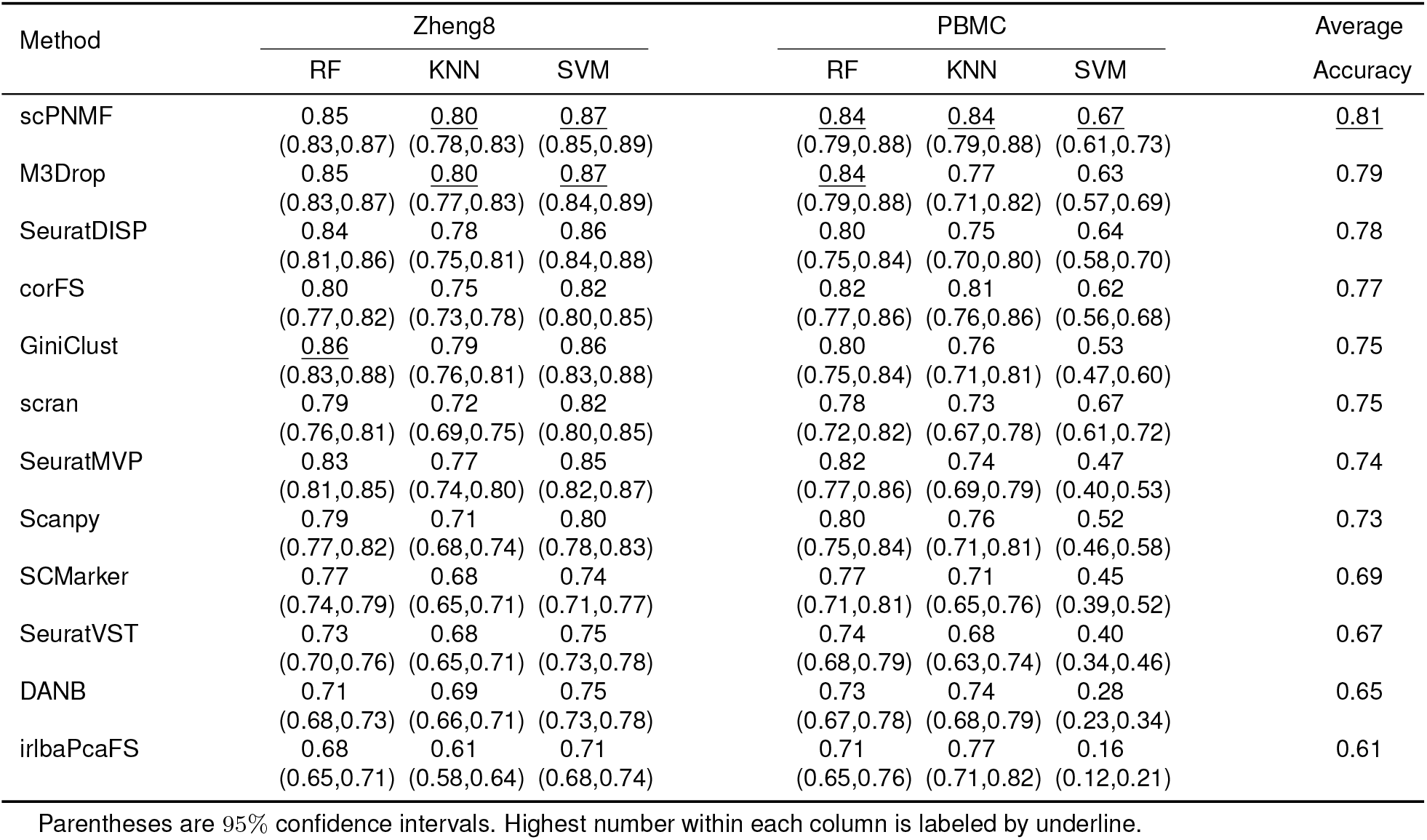
Prediction accuracy of cell types based on 100 informative genes selected by 12 gene selection methods in the two case studies with paired reference scRNA-seq data and targeted gene profiling data

## 4 Discussion

We propose scPNMF, an unsupervised gene selection and data projection method for scRNA-seq data. The major goal of scPNMF is to select a fixed number of informative genes to distinguish cell types and guide gene selection for targeted gene profiling experiments. Moreover, scPNMF can project a new targeted gene profiling dataset with the selected genes to the low-dimensional space that embeds a reference scRNA-seq dataset. We perform a comprehensive benchmark to evaluate scPNMF in terms of informative gene selection against the state-of-the-art gene selection methods. Our results show that scPNMF consistently outperforms existing methods for a wide range of informative gene numbers (from 20 to 500) on diverse scRNA-seq datasets. We also demonstrate that the informative genes selected by scPNMF can effectively guide gene selection for targeted gene profiling and lead to accurate cell type annotation on targeted gene profiling data based on reference scRNA-seq data.

Besides gene selection and data projection, scPNMF also works as a dimensionality reduction method with good interpretability. Each dimension in the low-dimensional space found by scPNMF can be considered as a new functional “feature” (as a linear combination of correlated and thus functionally related genes). Moreover, the mutual exclusiveness makes the PNMF bases used in scPNMF advantageous over the PCA bases in terms of removing confounding effects. For example, cell-cycle genes obscure the identification of cell types and should be removed from low-dimensional embeddings of cells. For PCA, cell-cycle genes affect many PCA bases, so the popular scRNA-seq pipeline Seurat implements a complicated approach that first calculates “cell-cycle scores” and then regresses each basis (principal component) on these scores to remove the effects of cell-cycle genes [12]. In contrast, cell-cycle genes are concentrated in only one PNMF basis, so it is easy to remove that basis to clear the effects of cell-cycle genes. Therefore, scPNMF has great potentials in deciphering cell heterogeneity in single-cell data by working as an interpretable dimensionality reduction method.

The current implementation of scPNMF focuses on single-cell gene expression data. Considering the rapid development of single-cell multi-omics technologies, we plan to extend scPNMF to accommodate other technologies that measure other genomics features such chromatin accessibility landscapes measured by single-cell ATAC-seq [42], or even to integrate data across multi-omics datasets. Another note is that the multimodality test for basis selection in scPNMF only accounts for discrete cell types but not continuous cell trajectories. Therefore, other tests or strategies are needed to select informative bases to capture biological variations along continuous cell trajectories.

An important question for gene selection is: how many genes should be selected as informative genes to fully capture the biological variations of interest? In our studies, we observe that, after the informative gene number reaches 200, the clustering accuracies based on the selected informative genes plateau for most gene selection methods including scPNMF. Therefore, 200 genes may be sufficient for capturing biological variations in scRNA-seq data. However, it remains challenging to decide the minimum number of informative genes, given that the underlying cell sub-population structure is data-specific and might be complex. We plan to explore this problem in future with the possible use of information theory.

## Software and code

The R package scPNMF is available at https://github.com/JSB-UCLA/scPNMF.

## Acknowledgements

We acknowledge the comments and feedback from the members of the Junction of Statistics and Biology at UCLA (http://jsb.ucla.edu).

## Supplementary Materials

### S1 Choice of parameters and robustness analysis

#### S1.1 Low rank K

In the development of scPNMF, motivated by the objective function of the PNMF method,

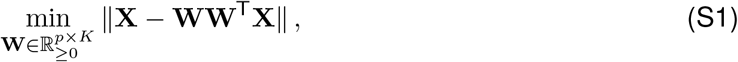

PNMF aims to inherit the advantages such as basis orthogonality and the ability to project the new data from PCA. However, a key constraint in PCA, **W**^T^**W** = **I**, is sacrificed in order to meet with the condition **W** *≥* 0 in PNMF. To get closer to PCA and thus attain its nice properties, we propose to use the normalized difference between **W**^T^**W** and **I** to measure the orthonality of **W**:

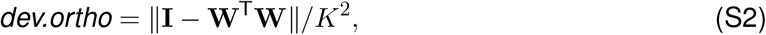

which is an implication of the performance in the downstream analysis as well.

It naturally follows a method for determining the number of basis, *K*: we perform PNMF for a sequence of *K*’s, calculate the *dev.ortho* measure for each 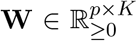 optimized by PNMF for each *K*, and then look at the plot of *dev.ortho* against *K*. Users can decide cutoff where it reaches stability or there is a clear elbow in the graph.

In Fig. S1, with Zheng4 [43] dataset, we demonstrate that (1) the *dev.ortho* measure is highly correlated with the performance of **W** in the downstream analysis; (2) in real data application, the *dev.ortho* measure shows a clear elbow pattern, which is helpful for users to determine *K*.

Empirically, we see that *dev.ortho* reaches stability at *K* = 20 for most scRNA-seq data. For the purpose of providing suggestion for users and saving computational energy, we set the default number of bases in scPNMF to be *K* = 20.

#### S1.2 R_0_: threshold for correlations between score vectors and cell library sizes in scPNMF step II: basis selection

In real data application, the threshold for correlations between score vectors and cell library sizes in scPNMF step II: basis selection, *R*_0_, needs to be pre-defined. In the field, researchers often use thresholds as accurate as with one decimal digit, such as 0.5. By empirically running K-means clustering on the seven datasets (see Table S3) with different thresholds *{*0.5, 0.6, 0.7, 0.8, 0.9*}*, as shown in Fig. S2, we suggest setting *R*_0_ = 0.7 for *K ≥* 10, and more conservatively, *R*_0_ = 0.8 when the basis number *K* is small (*K <* 10).

### S2 Functional annotation

We use the R package clusterProfiler [Y] to perform GO analysis. We set the gene ontology as “BP”, adjusted *p*-value cutoff as 0.1. The output GO terms are simplified by clusterProfiler.

In this paper, we only perform a very conservative filtering based on functionality. We define the common housekeeping gene list as *ACTB, ACTG1, B2M, GAPDH, MALAT1*. If the top 10 high weight genes from one basis contain any of these genes, this basis will be filtered out.

### S3 Data preprocessing

scPNMF only performs minimum data preprocessing to avoid information loss. Denote a scRNA-seq count matrix scPNMF further investigates as **X**^*C*^ *∈* N^*p×n*^, with rows representing *p* genes and columns representing *n* cells. Users make the log count matrix 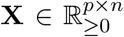 by taking the log transformation with a pseudo count 1:

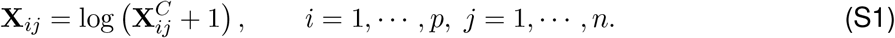

scPNMF takes the log count matrix 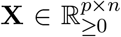 as the input. With log transformation, the effect of a few extremely large counts will be alleviated, and the transformed continuous values are more flexible to model. We introduce the pseudo count 1 to avoid negative and infinite values in the later PNMF optimization step.

For scRNA-seq data used in this paper (Table S1), we filtered out genes that are expressed in fewer than 5% of the cells, and then filtered out cells that are expressed in fewer than 5% of the remaining genes. Additionally, *MALAT1*, mitochondrial and ribosomal genes are filtered for datasets PBMC10x and PBMCSmartSeq according to the reference paper [44]. Users are able to adjust the filtering process before they input the log count matrix into scPNMF.

### S4 Details in informative gene selection and clustering

In this paper, we compare scPNMF with other 11 different informative gene selection methods (Table S2). Some gene selection methods cannot let users pre-define an arbitrary gene number; for those methods (e.g., SCMarker [16]), we shift the tuning parameters until their output gene numbers equals the desired gene number. Therefore, their outputs might not achieve their the optimal results.

We apply three clustering algorithm, Louvain clustering (by Seurat), K-means clustering (by R function kmeans), hierarchical clustering (by R function hclust). We perform PCA on informative genes and use the top 20 PCs for clustering. The Adjusted Rank Index (ARI) is as the metric of clustering accuracy. ARI is defined as:

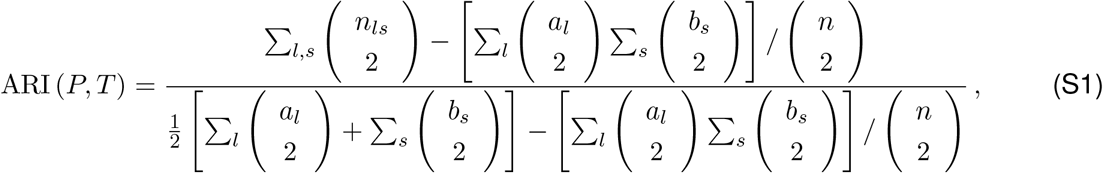

where *P* = (*p*_1_, *· · ·, p*_*l*_) denotes the inferred cluster labels, and *T* = (*t*_1_, *· · ·, t*_*s*_) denotes the true cluster labels. *l* and *s* are not necessarily to be equal. 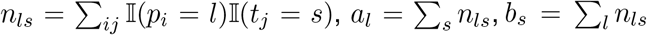 ARI *∈* [0, 1], an ARI value close to 1 means more accurate inferred clusters. To minimize the effects caused by parameters (resolution *r* in Louvain and number of cluster *k* in K-means and hierarchical clustering), we try a sequence of parameters:

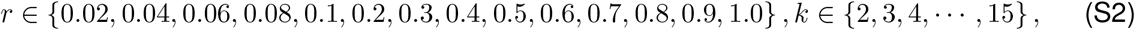

and use the average of top three high ARI across different parameters as the final output.

### S5 Details in new data projection and cell type prediction

We use two datasets, Zheng8 and PBMC10x, as the reference scRNA-seq datasets. For Zheng8 dataset, we first use scDesign2 [39] to learn the underlying parameters, and then simulate a new dataset with same genes and cell types but 100 times higher sequencing depth compared to the Zheng8 dataset. For PBMC10x dataset, we use the PBMCSmartSeq dataset, which measures the exact same example and contains all genes measured in PBMC10x. Given *M* selected genes, the simulated Zheng8 and PBMC10x are extracted with those certain genes, and play role as the “pseudo” targeted gene profiling only measuring *M* genes.

For cell type prediction, we project every targeted gene profiling dataset and its scRNA-seq reference on the same low-dimensional space, which mainly follows the idea from scPred [45]. When applying scPNMF, we use the weight matrix **W**_*S*,(*M*)_ to project both the reference dataset and the targeted gene profiling dataset. For other gene selection methods, we first subset the reference dataset with only *M* selected genes, run PCA to get a weight matrix **W**_PCA_, and use it to project both the reference dataset (with only *M* genes) and targeted gene profiling dataset. After getting two low-dimensional embeddings of reference and targeted gene profiling data, we run the Harmony algorithm [32] to remove the technical variations between two low-dimensional embeddings. Then we apply three classification algorithms, random forest (`rf`), k-nearest neighbors (`knn`) and support vector machine with radial kernel (`svmRadial`) in R package caret [K]. When fitting the training model, we use 5-fold cross-validation with three repeats.

**Table S1:** Top 10 high weight genes in each PNMF basis of FretagGold dataset

| Basis | Gene symbol | Description |
| --- | --- | --- |
| 1 | *RPS2, TMSB4X, GAPDH, RPL41, RPL13, FTH1, MALAT1, COX2, RPL10, RPS18* | Highly expressed housekeeping genes |
| 2 | *CD74, PTGR1, HLA-B, ALDH3A1, C15orf48, LCN2, IGFBP3, SAA1, CXCL1, HLA-DRA* | Immune-related genes |
| 3 | *SEC61G, CDK4, CCN1, G0S2, ELOC, VOPP1, EGFR, F3, CDKN2A, EPCAM* | Tumor-related genes (oncogenes, tumor suppressor genes) |
| 4 | *H4C3, CKS1B, HMGB2, SMC4, PTTG1, KPNA2, CCNB1, CDKN3, CKS2, CDC20* | Genes related to mitotic cell cycle |
| 5 | *HSPB1, UBE2S, CALD1, TMEM256, FIS1, ISOC2, ZN-HIT1, C20orf27, NDUFA3, PPP2R1A* | Genes related to mitochondrion |

**Table S2:**
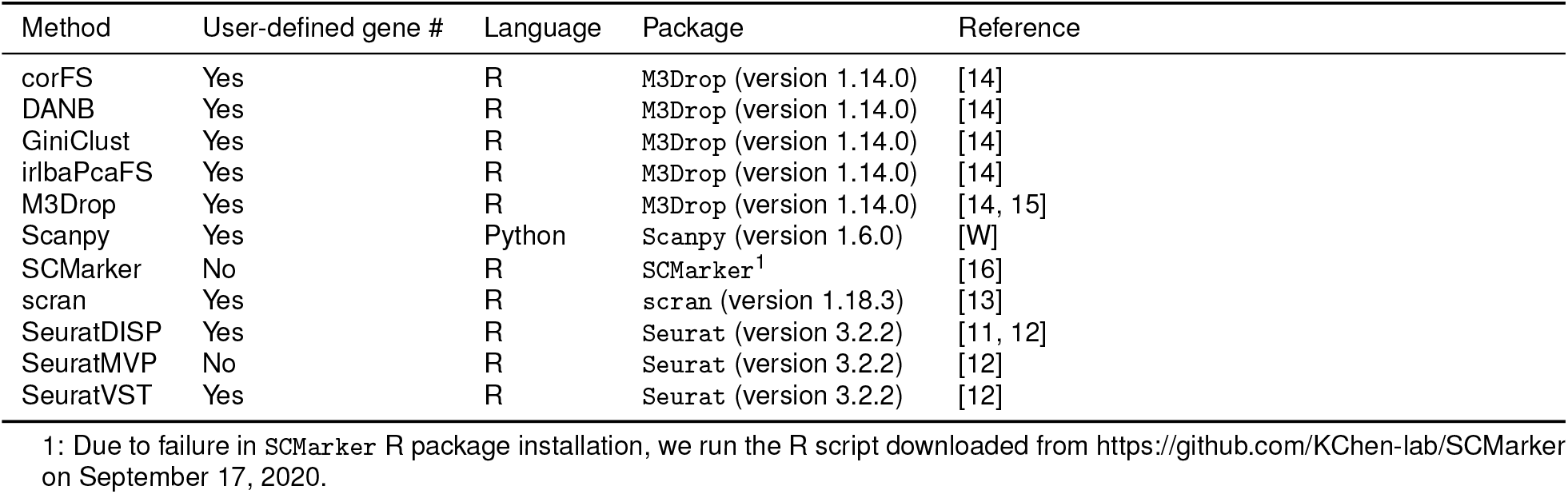
Overview of informative gene selection used in this study

**Table S3:**
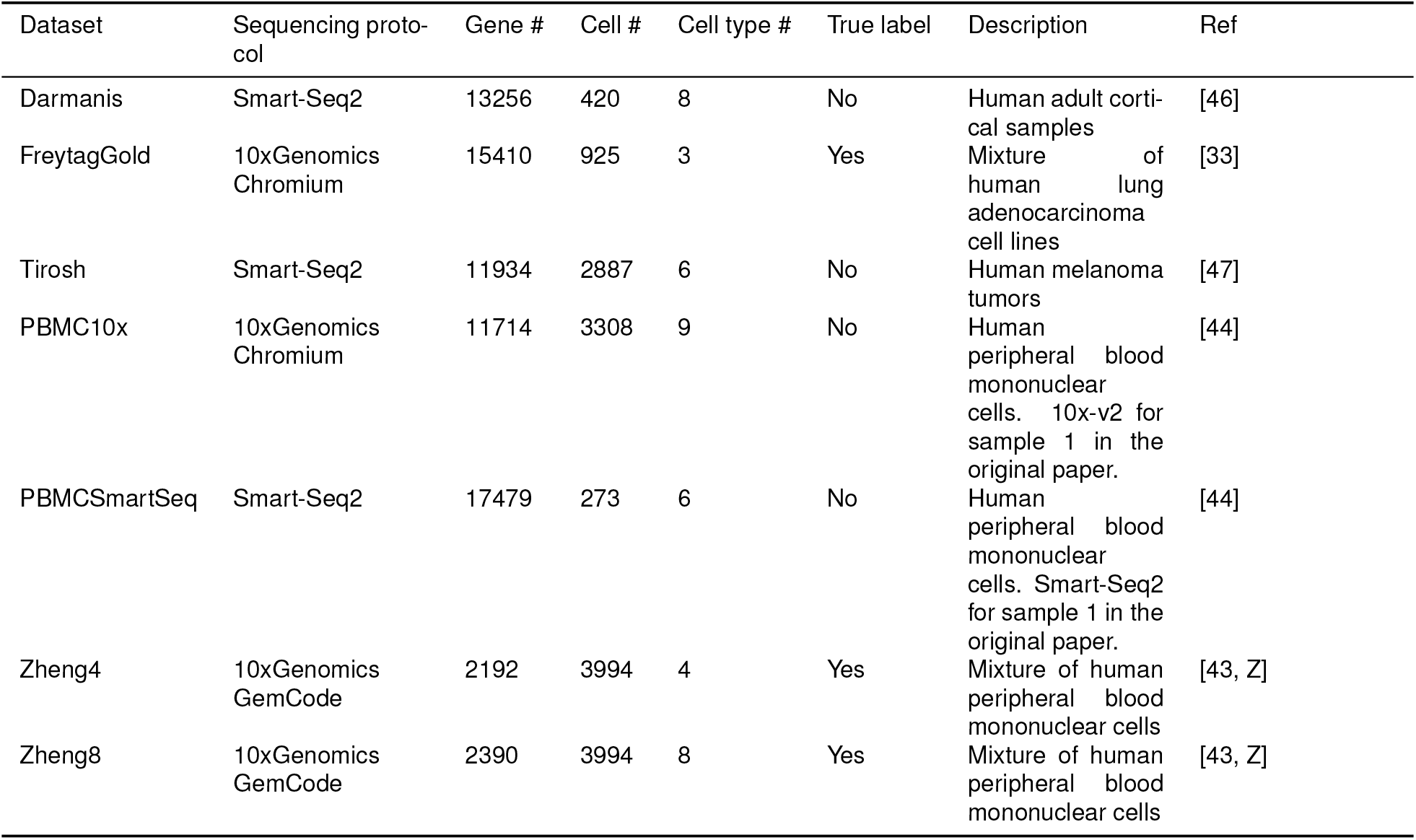
Overview of datasets used in this study

| Dataset | Sequencing protocol | Gene # | Cell # | Cell type # | True label | Description | Ref |
| --- | --- | --- | --- | --- | --- | --- | --- |
| Darmanis | Smart-Seq2 | 13256 | 420 | 8 | No | Human adult cortical samples | [46] |
| FreytagGold | 10xGenomics Chromium | 15410 | 925 | 3 | Yes | Mixture of human lung adenocarcinoma cell lines | [33] |
| Tirosh | Smart-Seq2 | 11934 | 2887 | 6 | No | Human melanoma tumors | [47] |
| PBMC10x | 10xGenomics Chromium | 11714 | 3308 | 9 | No | Human peripheral blood mononuclear cells. 10x-v2 for sample 1 in the original paper. | [44] |
| PBMCSmartSeq | Smart-Seq2 | 17479 | 273 | 6 | No | Human peripheral blood mononuclear cells. Smart-Seq2 for sample 1 in the original paper. | [44] |
| Zheng4 | 10xGenomics GemCode | 2192 | 3994 | 4 | Yes | Mixture of human peripheral blood mononuclear cells | [43, Z] |
| Zheng8 | 10xGenomics GemCode | 2390 | 3994 | 8 | Yes | Mixture of human peripheral blood mononuclear cells | [43, Z] |

**Figure S1:**
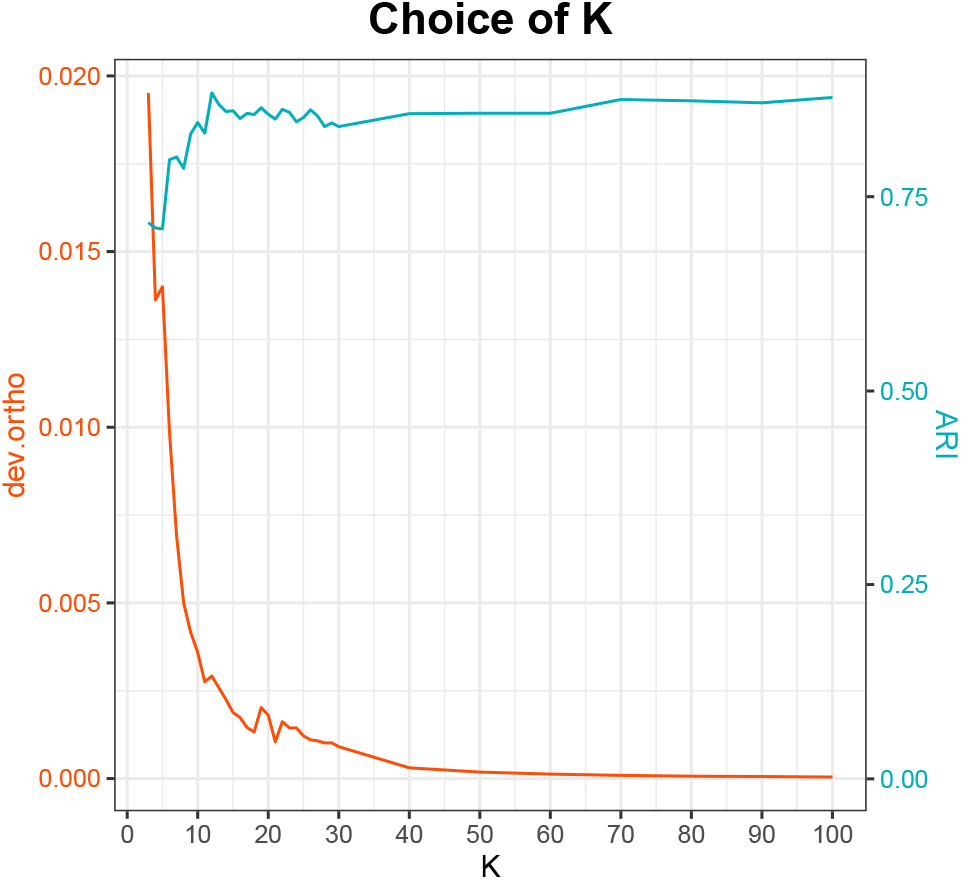
Comparison of *dev.ortho* and K-means ARI against low rank *K* on Zheng4 [43] dataset.

**Figure S2:**
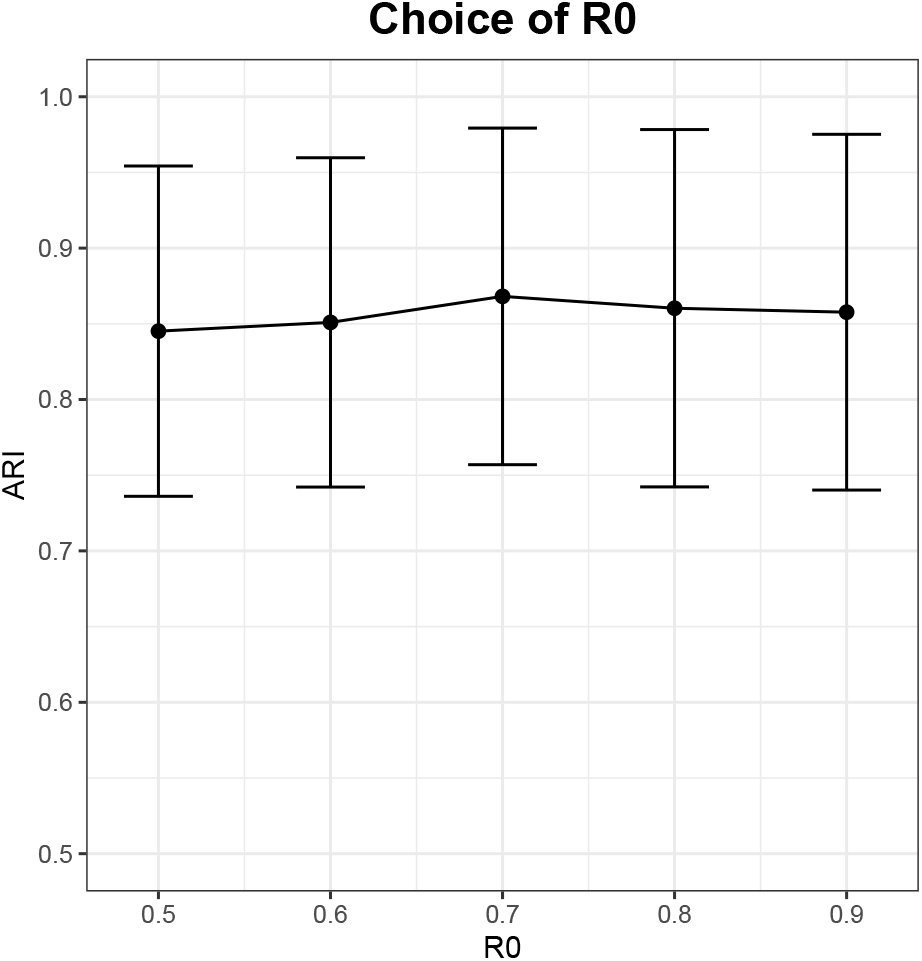
Comparison of K-means ARI against *R*_0_, the threshold for correlations between score vectors and cell library sizes in scPNMF step II: basis selection. The mean ARI and the error bars are calculated across seven datasets (See Table S3).

**Figure S3:**
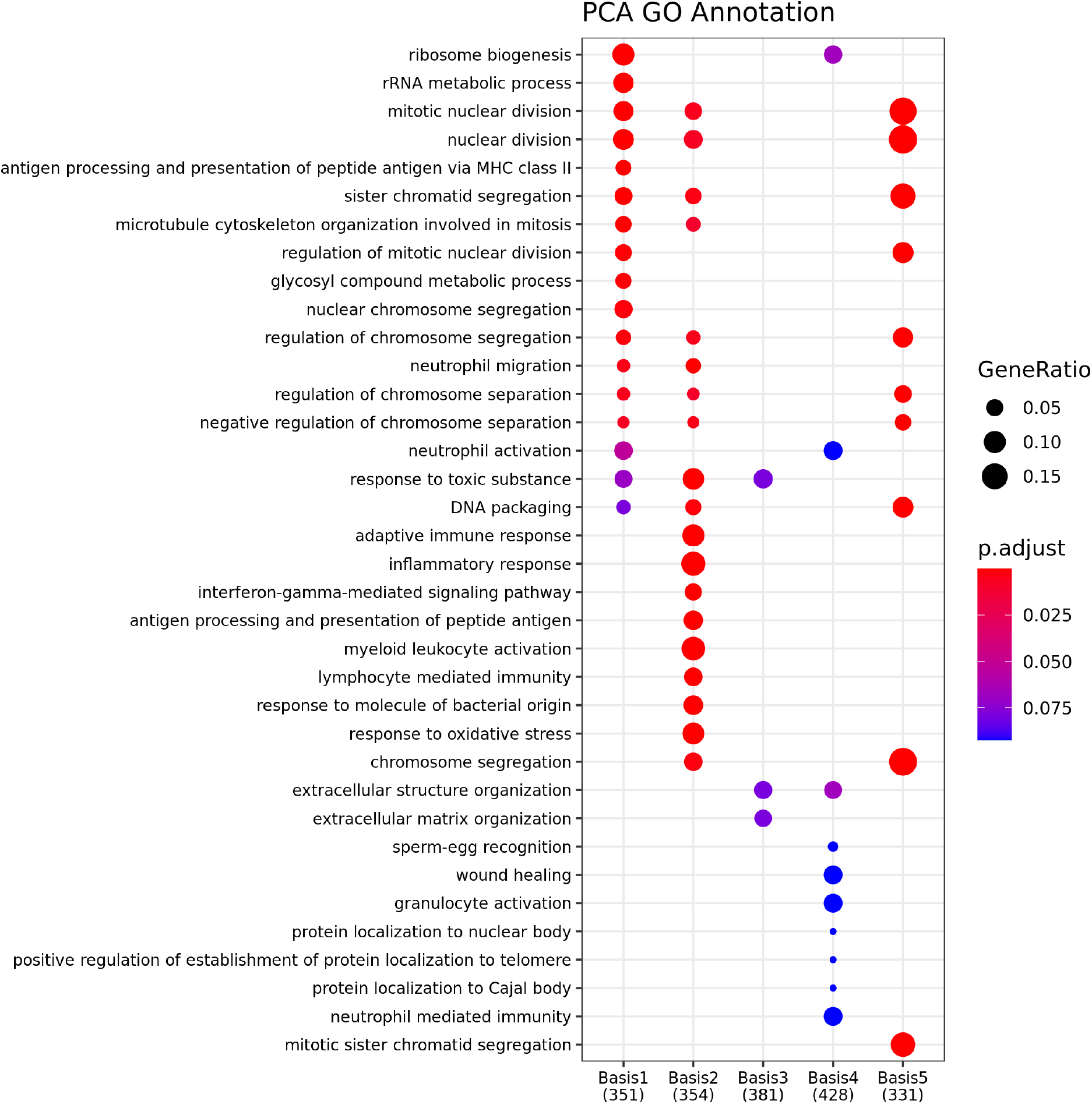
GO annotation on weight matrix of PCA. The enriched GO terms between basis are largely overlapped.

**Figure S4:**
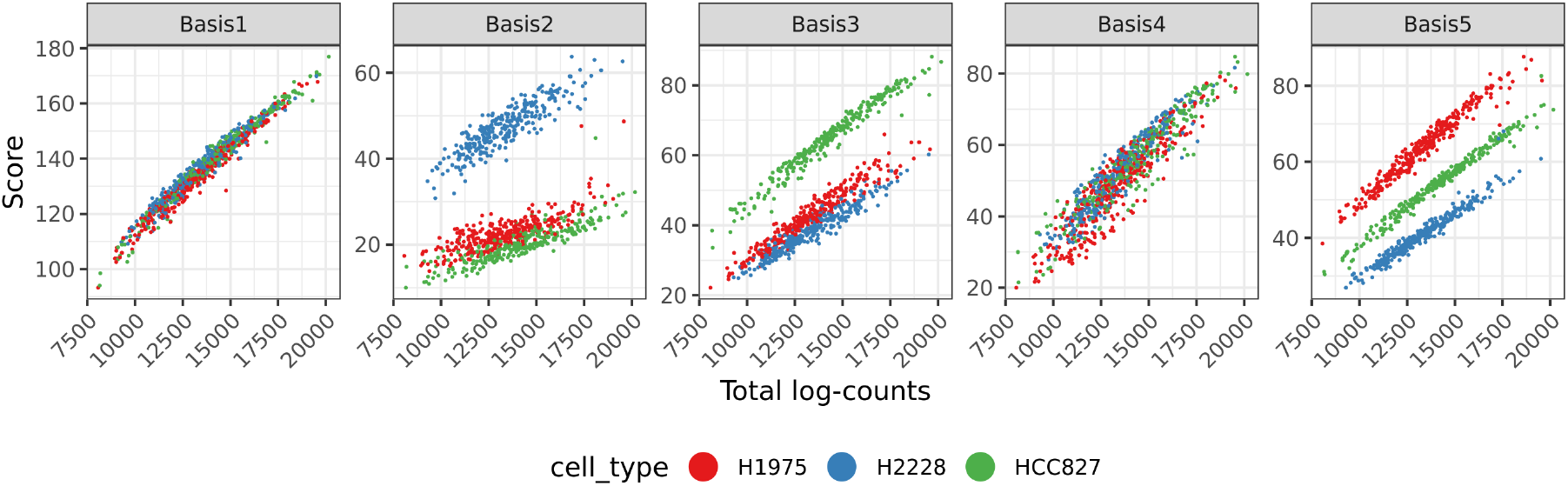
scPNMF scores versus total log-counts of FregGold dataset colored by cell types. Basis 2 distinguishes H2228 from the other two cell types and basis 3 distinguishes HCC827 from the other two cell types.

**Figure S5:**
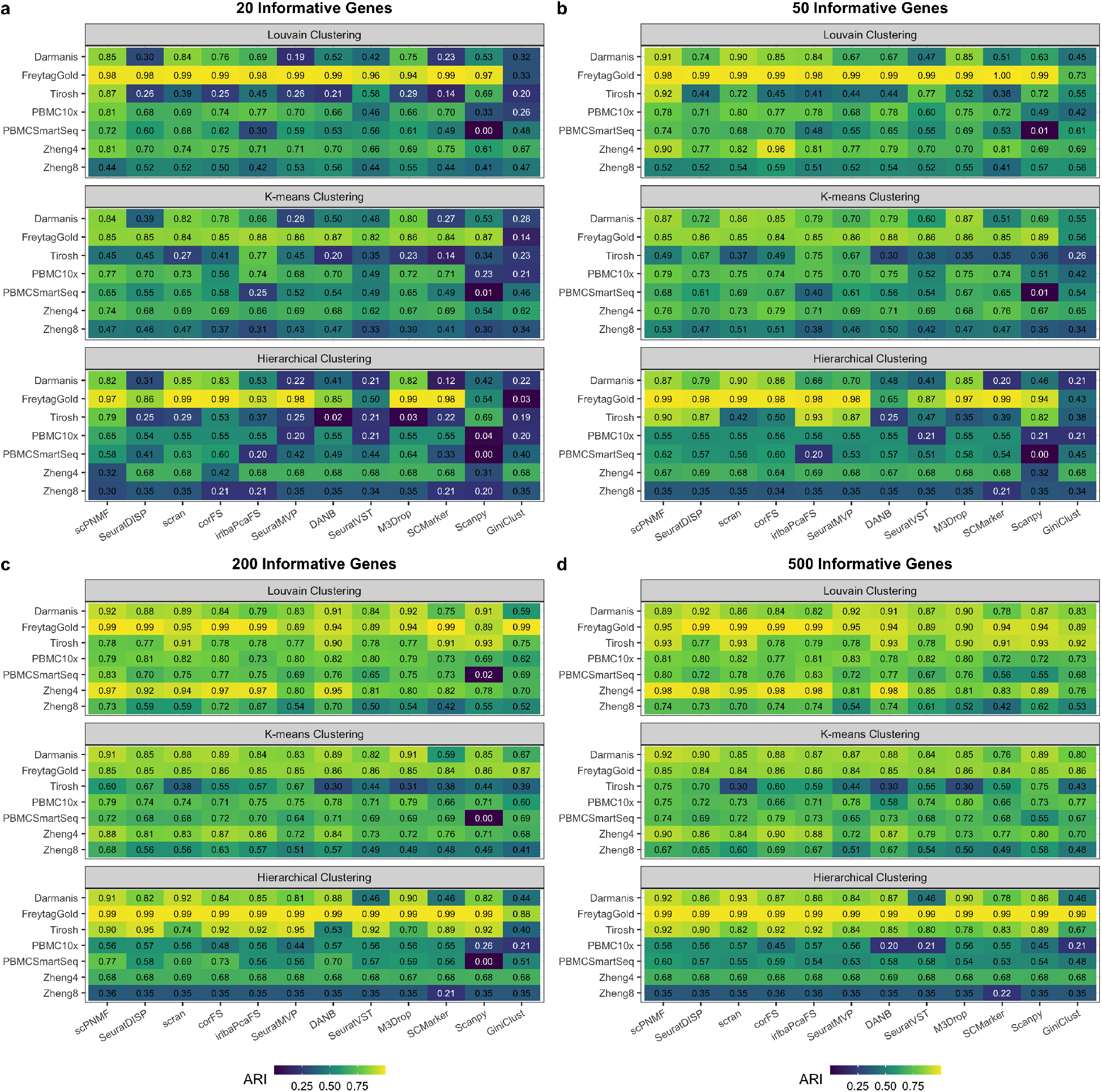
Benchmarking scPNMF and other informative gene selction methods using 20, 50, 200, 500 genes.

**Figure S6:**
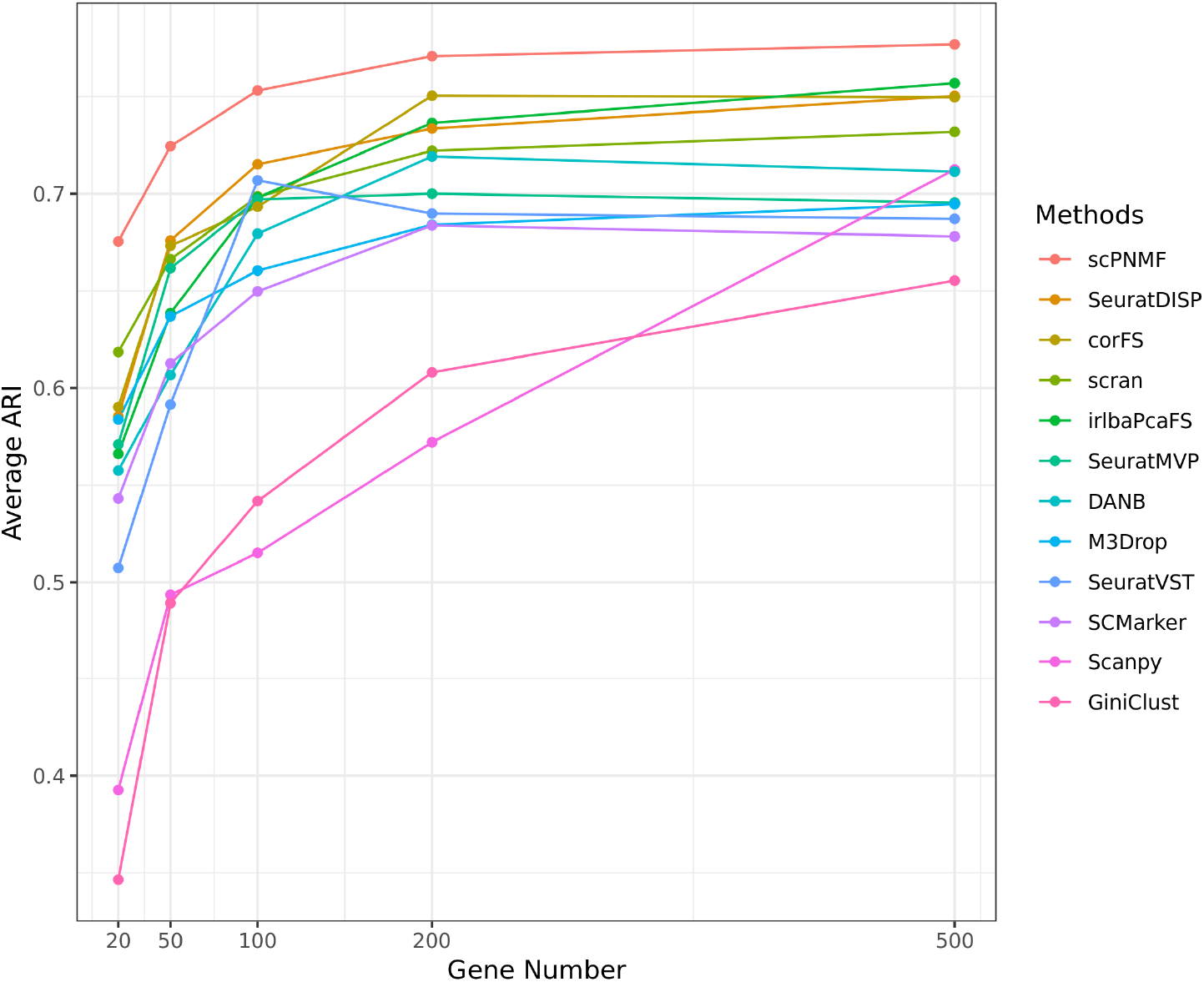
Comparison of overall average ARI of different methods versus gene numbers. The *y*-axis indicates the average ARI values across seven datasets and three clustering methods for each gene selection methods.

